# THE SUBTHALAMIC NUCLEUS IS INVOLVED IN SOCIAL RECOGNITION MEMORY IN RATS

**DOI:** 10.1101/2023.02.07.527559

**Authors:** Cassandre Vielle, Nicolas Maurice, Florence Pelletier, Emilie Pecchi, Christelle Baunez

## Abstract

Human social behavior is a complex construct requiring a wide range of cognitive abilities and is critically impaired in numerous neuropsychiatric diseases. Living in complex social groups, rodents offer suitable models to elucidate neural processing of social cognition. Recently, a potential involvement of the subthalamic nucleus (STN) in rats’ social behavior has been pointed out. For example, we showed that STN lesions abolish the modulatory effect of the familiarity on the rewarding value of social stimuli, questioning the involvement of STN in peer recognition. In this study, we thus assess the effects of STN lesions and optogenetic manipulations on peer and object recognition. STN optogenetic inhibition, like lesions, impair social recognition memory, while STN optogenetic high-frequency (HF) stimulation leads to a specific alteration of social encoding memory. None of these manipulations seem to interfere with social investigation, objects recognition memory, nor social novelty preference. Finally, STN optogenetic inhibition, but neither HF-stimulation, nor lesions, leads to an alteration of the cage-mate recognition memory. Overall, these results show that physiological activity of STN is necessary for rats to show a proper social recognition memory performance and question the possible detrimental effects of STN deep brain stimulation on these processes in human patients.

## Introduction

Like humans, rats live in a complex social environment (Barnett, 1967), requiring a large panel of social abilities, from simply categorizing others (e.g., male *vs*. female or kin *vs*. stranger) or reading basic emotions, to complex social learning (i.e., the ability to learn from others’ actions), helping behavior or cooperative behaviors (for review, see Schweinfurth, 2020). Among all these social skills, social recognition memory refers to the ability to recognize the identity of a peer based on previous encounter. Likewise, during an initial encounter, a rat investigates a stranger peer, collecting information on the identity of this novel conspecific and storing a social memory. During a second encounter with the same individual, the rat will retrieve the social information about this peer, which enables engagement of appropriate interactions (e.g., approach or avoidance, mating, attack) with this ‘now familiar’ conspecific. Thus, social recognition is a fundamental process in gregarious species, allowing complex social behaviors.

Since laboratory rats show a preference for novel stimuli, their social recognition abilities may be assessed by scoring their investigatory behavior toward a novel vs familiar peer (Engelmann et al., 1995; Thor & Holloway, 1982). Yet, it is noteworthy that deficits in memory, approach, attentional processes or reward processing may also affect animal’s investigatory behavior. The use of these tests and others in laboratory rodents allowed to identify some of the specific neurobiological substrates of social recognition memory in rodents, comprising the olfactory bulb (Richter, 2005), the hippocampus (Chai et al., 2021; Hitti & Siegelbaum, 2014), the medial (Ferguson et al., 2001) and basolateral amygdala (Wang et al., 2014) and some cortical areas, such as the anterior cingulate (Rudebeck et al., 2007) and the prefrontal (Tan et al., 2019) cortices. Interestingly, some of these structures seem to be involved specifically in sub-processes of social recognition memory (i.e., acquisition, maintenance or retrieval of social recognition memory) (Kogan et al., 2000; Richter, 2005). For instance, inhibition of the medial amygdala before retrieval, but not learning, session impairs social recognition memory in mice (Noack et al., 2015).

Historically known as the glutamatergic relay of the basal ganglia, merely acting on motor functions, the subthalamic nucleus (STN) has never been studied for its involvement in social recognition memory. Nevertheless, basal ganglia also form associative and limbic loops with the cortex and the thalamus (Alexander et al., 1986). At the interface of these associative and limbic systems, the STN receives cortical projections from a wide variety of regions, such as the anterior cingulate and the prefrontal cortices and sends projections to most components of the basal ganglia. Furthermore, segregation of STN afferent and efferent projections enables the identification of at least 3 distinct functional territories on this nucleus: the motor dorsolateral STN, the associative medial STN and the limbic ventral STN (Hamani, 2004). This anatomical organization suggests that the STN has actually a central position in basal ganglia circuitry (Parent & Hazrati, 1995) and is involved in limbic and cognitive functions of this circuitry (Maurice et al., 1998). Indeed, behavioral studies have shown that the STN is involved in a large panel of high level cognitive and emotional functions (for review, see Baunez & Lardeux, 2011).

Regarding the role of the STN on social behavior, Tan et al. (2011) showed that electric high frequency (HF) stimulation of the STN increases sniffing behavior in Parkinsonian-like rats during social interaction tests, while Reymann et al. (2013) found that STN lesions decreased some social behaviors (i.e., investigation, contact and tail manipulation) in control rats. Finally, our group showed that STN lesions blunt the modulatory effect of the familiarity on the rewarding value of social stimuli. In control rats, but not in STN-lesioned ones, the presence of a stranger peer is more rewarding, and decreases further cocaine intake, than the presence of a familiar one (Giorla et al., 2022). Similarly, in control rats, but not in STN-lesioned animals, the playback of positive ultrasonic vocalizations (USV) emitted by a stranger rat is more reinforcing and further decreases cocaine consumption, than if emitted by the cage-mate (Vielle et al., 2021). Altogether, these results suggest that the STN could be involved in social recognition.

This study aimed thus at exploring the involvement of STN in social recognition. To better characterize its possible role, we used lesions and optogenetic tools to either inhibit or stimulate at high frequency (130Hz) the activity of STN neurons. We assessed the effects of these STN neuromodulations on rats’ ability to discriminate a stranger from a previously encountered peer, immediately after the first encounter, or after a 30-mn intertrial delay. We also tested the effects of STN manipulations on the cage-mate recognition to better the extent of the social recognition memory alteration, and object recognition memory, to check the specificity of this alteration. Finally, we controlled the effect of both optogenetic manipulations activity, using *in vivo* recordings of STN activity in anesthetized animals. Overall, the mixed lesional and optogenetic approach allowed us to differentiate the effects of short- and long-term manipulation of STN activity on social recognition. Likewise, we were able to target a possible involvement of the STN in a sub-process of social recognition (i.e., encoding and recall of social memory) with optogenetics tools, and a possible compensatory effect of its lesion. Our results may have critical consequences for the follow-up of patients treated with STN deep brain stimulation (DBS) for Parkinson’s Disease (PD) or obsessive-compulsive disorders (OCD).

## Material and methods

### Animals

55 male Lister Hooded rats (Charles River Laboratories, Saint-Germain-sur-l’Arbresle, France) weighing ∼400 g were used in this study. Only male rats were used, because social interaction is more rewarding for them than for females (Douglas et al., 2004). Animals were housed in pairs in temperature-controlled room, maintained under a 12 h inverted light/dark cycle, with unlimited access to water and food (Scientific Animal Food and Engineering, Augy, France). Animals were handled daily for habituation. All experiments were conducted during the dark cycle (7am-7pm). All animal care and use conformed to the French regulation (Decree 2010-118) and were approved by the local ethic committee and the French Ministry of Agriculture under the License #3129.01 and followed the 3R European rules.

### Virus

To transfect the STN neurons, we used AAV5 with the recombinant protein expression under CamKIIa-promoter control, obtained from UNC Vector Core (Chapel Hill, USA). The control, inhibition and HF-stimulation groups received respectively the following AAV5 constructs: AAV5-CaMKII-EYFP, AAV5-CaMKII-ArchT3.0-p2A-EYFP-WPRE and AAV5-CaMKII-hChR2(E123T/T159C)-p2A-EYFP-WPRE.

### Optogenetic manipulation

Homemade optic fibers (230µm, Thorlabs) were connected, via an optic coupler (FCMM625-50A, Thorlabs), to a 200mW 532nm DPSS laser and light pulses were generated under the control of a signal generator (DS8000 Digital Stimulator, World Precision Instruments) with 2ms light pulse at 130Hz pulse train for STN stimulation and 15s light pulse at 0.2Hz pulse train for STN inhibition. The laser power at fiber tip was set, using a power meter (PM20A, Thorlabs), at 10mW for STN HF-stimulation and 5kW for its inhibition. The effect of these light delivery parameters on STN neurons firing pattern were previously tested *in vitro* and *in vivo* (Vielle et al, 2023). Nevertheless, we controlled here the effects of STN photo-inhibition and HF photo-stimulation, via extracellular recordings of STN activity in anesthetized animals (see above).

### Surgeries

All rats were anesthetized with ketamine (Imalgene, Merial, 100 mg/kg, *i*.*p*.) and medetomidine (Domitor, Janssen 30 mg/kg, *i*.*p*.), which was reversed at the end of the surgical procedure with an injection of atipamezole (Antisedan, Janssen, 0.15 mg/kg, *i*.*m*.). They received an antibiotic treatment with amoxicillin (Duphamox, LA, Pfizer, 100 mg/kg, *s*.*c*.) and were administered meloxicam (Metacam, Boehringer Ingelheim, 1 mg/kg, *s*.*c*.) for analgesia.

Animals were placed in the stereotaxic frame (David Kopf apparatus) and receive 0.5µL bilateral injections of virus (for optogenetic rats) or 53 mM ibotenic acid (9.4 mg/mL, for STN-lesioned rats) or vehicle solution (phosphate buffer, 0.1 M; for sham-control rats) into the STN (with tooth bar set at - mm), at the following coordinates : anterior/posterior = −3.7 mm ; lateral = ±2.4 mm from bregma ; dorsoventral = −8.4 mm from skull; from Paxinos & Watson, 2007. For optogenetic rats subjected to behavioral testing, optic fibers were implanted 0.5 mm above each injection site and fixed on the skull embedded in a head-cap made of dental cement. These rats were habituated to be connected to the optic coupler several days before the experiments. Optogenetic rats were allowed 30 days for recovery and satisfying expression of the opsins. Sham-control and STN-lesioned rats were allowed 10 days for recovery.

#### A) Behavioral procedures

To assess the effects of STN lesions on social recognition, 6 Sham-control and 9 STN-lesioned rats were subjected to a control objects discrimination test, a social habituation-dishabituation test and various social discrimination tests: 1) with no delay, 2) with a 30-min delay between the two encounters, and 3) with the cage-mate.

To test the possible effect of a short-term modulation of the STN on the subprocesses of social recognition, 30 optogenetic rats (10 for EYFP-control, 10 for STN inhibition and 10 for STN HF-stimulation groups) were subjected to social (with no delay and with 30-min delay) and objects discrimination tests twice (one time with the laser ON during the first encounter and a second time with the laser ON during the 2^nd^ encounter). We also tested their ability to recognize their cage-mate, with the laser ON during stimuli encounter.

To reduce the number of animals used, the same cohort served for various procedures. Every rat was thus subjected to each test, in a counterbalanced order. For each individual, a waiting period of at least one day occurred between each test as a subject, since repeated daily testing does not affect social investigation (Thor & Holloway, 1982), or one week as a stimulus for a test before being the subject of a test.

### Apparatus

All tests took place in a rectangular plastic arena (40 cm length, 40 cm high, 30 cm depth), with fresh litter, changed between each test. Arenas and objects were cleaned between each test with a solution of hydrogen peroxide. The experimental room was quiet and dimly illuminated. Subject animals were habituated to the arena during 30-min before each test. Stimulus rats had approximatively the same age and weight than the subject rat and were habituated to be handled. Stimulus rats and objects were different for every test condition. Their order of presentation (1^st^ and 2^nd^ or only 2^nd^ encounter) to the subject rat was counterbalanced between animals. During each encounter, the subject animal was free to explore the stimuli. Investigation time was defined, as previously described in Engelmann (1995), as the time spent by the subject to actively explore (sniffing with the tip of the nose within approximately 1 cm, following, nosing, grooming, licking) a stimulus (rat or object). Every encounter of the subject rat with a stimulus was recorded with a webcam connected to a computer, using Bonsai (Open Ephys) and investigatory behavior was scored by a trained observer. For each encounter, if the total investigation time was less than 30-sec, or if rats fought, the results of the subject rat for this test were discarded from the data analyses.

#### 1) Social habituation-dishabituation test

To assess a possible effect of the STN lesion on rats’ ability to recognize a peer, sham control and STN-lesioned rats were subjected to an habituation-dishabituation procedure (Winslow & Camacho, 1995). In this procedure, the subject rat was exposed four times to the same unfamiliar stimulus rat. These 5-min encounters were interleaved with 10-min intertrial intervals, during which the subject remained in the arena, while the stimulus was replaced in its home-cage. These four sessions constituted an “habituation phase”, during which the exploratory behavior of the subject is supposed to decrease across sessions, since the stimulus rat become familiar. After the 4^th^ intertrial interval, the subject rat was exposed to a novel unfamiliar peer for 5-min. This encounter constitutes the “dishabituation phase”, during which the exploratory behavior of the subject is supposed to reinstate. This last session was used to control fatigue or disinterest of the subject rat.

#### 2) Social discrimination test with no delay, or with 30-min intertrial delay

To confirm and better characterize the effect of the STN lesion on social recognition, sham-control and STN-lesioned rats were subjected to a social discrimination test (Engelmann et al., 1995, 2011). This procedure consisted in 2 encounters. During the first encounter (encoding phase), the subject rat was exposed for 5-min to an unfamiliar peer. Then, immediately after or after a 30-min intertrial interval (respectively social discrimination test with no delay, and with 30-min delay), during a second 5-min encounter (retrieval phase), the subject rat was exposed to two peers: the previously encountered one, which was now ‘familiar’, and, simultaneously, a novel unfamiliar peer. During this session, considering rats’ preference for social novelty, the subject was expected to spend more time exploring the novel peer, than the previously met one.

To test the possible effect of a short-term modulation of STN activity on a subprocess of social recognition (i.e., encoding and recall), optogenetic rats were also subjected twice to these 2 social discrimination tests. The first time, the laser activation was ON during the 5-min of the first encounter, but OFF during the 5-min of the second encounter, and *vice versa* the second time. The order of presentation for these 2 laser activation conditions was pseudo-randomly applied, so that every rat was tested in each condition, in a counterbalanced order. To reduce any bias due to the connection to the optic fibers, we connected the rats’ optic fibers to the optic coupler, whatever the laser activation ON or OFF.

#### 3) Object discrimination test with 30-min delay

To assess the specificity of the effect of STN modulation on social recognition memory, we adapted the social discrimination test with 30-min delay to test rats’ ability to discriminate the familiarity of a non-social stimulus. Then, this procedure was the same than for the social discrimination test with 30-min delay except that stimulus rats were replaced by objects. The two objects used in this experiment had approximatively the same size and were made with the same material.

#### 4) Cage-mate recognition test

To test the extent of the social memory recognition alteration due to STN neuromodulation, sham-control, STN-lesioned and optogenetic rats were subjected to a cage-mate recognition test. In this test, the subject rat was exposed, simultaneously, to its cage-mate and a novel unfamiliar rat, for a single 5-min encounter. Of note, one STN-lesioned rat was excluded from the results because its cage-mate died before the test.

#### *B)* In vivo *extracellular recordings of STN activity*

*In vivo* electrophysiological recordings were performed in n=3 and n=7 rats to assess the effect of respectively STN optogenetic inhibition and HF-stimulation. Briefly, rats were anesthetized with a mixture of ketamine/xylazine (100/10 mg/kg, *i*.*p*.) supplemented as needed during the recording session.

They were mounted in a stereotaxic head frame (Horsley-Clarke apparatus; Unimécanique, Epinay-sur-Seine, France). Their body temperature was maintained at 36.5°C with a homeothermic blanket controlled by a rectal probe (Harvard Apparatus, Holliston, MA). Single-unit activity of neurons in the STN was recorded extracellularly using glass micropipettes (25-35 MΩ) filled with a 0.5 M sodium chloride solution. Action potentials were recorded using the active bridge mode of an Axoclamp-2B amplifier (Molecular Devices, San Jose, CA), amplified, and filtered with an AC/DC amplifier (DAM 50; World Precision Instruments). Data were sampled on-line at 10 kHz rate on a computer connected to a CED 1401 interface and off-line analyzed using Spike2 software (Cambridge Electronic Design, Cambridge, UK). Optical fiber (230 μm-Thorlabs) was inserted with a 15° angle and the fiber tip was lowered just above the STN and connected to a a 200mW 532nm DPSS laser. The entry point had the following coordinates: AP: -3.7 mm, ML: +1.0 mm. The fiber tip was at a depth of 7.3 mm from the cortical surface.

The pattern of STN neurons activity was analyzed at least 20s before, 180s during and 20s after photo-modulation. The parameters of light stimulation were the same than those used in behavioral experiments.

### Immuno-histology

At the end of the experiments, sham-control and STN-lesioned rats were deeply anesthetized with isoflurane and then euthanized using an intracardiac injection of pentobarbital sodium (Dolethal, Vetoquinol, 200mg/kg). Optogenetic rats were anesthetized with an overdose of a cocktail of ketamine and medetomidine (Imalgene, Merial and Domitor, Janssen 30 mg/kg, *i*.*p*.) and transcardially perfused with 0.1M PBS followed by 4% paraformaldehyde dissolved in PBS. Brains were removed and, after a cryoprotection in 30% sucrose for optogenetic brains, frozen in isopentane (Sigma-Aldrich) and kept at −80°C before being coronally sectioned at 40 µm thickness, using a cryostat.

Brain sections from sham-control and STN-lesioned rats were stained with thionine (Sigma-Aldrich) to assess the location and extent of STN lesions, characterized by neuronal loss and associated gliosis. Brain sections from EYFP-control and STN optogenetic inhibition rats were examined for optic fibers location and for native fluorescence expression with an epifluorescence microscope (Zeiss, Imager.z2) immediately after being cut. In brain sections from STN HF-stimulation animals, native fluorescence was present but required immuno-staining to control the exact boundaries of the viral expression. After PBS rehydration (3 × 5 min), brain sections underwent a 90 min permeation step (PBS, 1% bovine serum albumin (BSA) 2% normal goat serum (NGS), 0.4% TritonX-100), PBS washes (3 × 5 min) and incubation with primary antibody (mouse anti-GFP, A11120, Life technologies; 1:200, in PBS 1% BSA, 2% NGS, 0.2% TritonX-100) at 4°C overnight. Sections were then washed with PBS (3 × 5 min), followed by 2h incubation at room temperature with secondary antibody (Goat anti-mouse Alexa 488, A11011, Life technologies, 1:400 in PBS 1% BSA, 2% NGS) and finally washed with PBS (3 × 5 min).

### Statistical analysis

All variables were expressed as mean number ± SEM and p-value threshold was set at α=0.05. Analyses and graphs were carried out using Prism (GraphPad). For the social habituation-dishabituation test, we compared the time spent by the subject rat to explore a conspecific between each session using Friedman test, followed by Dunn multiple comparisons for post hoc analysis, separately for sham-control and STN-lesioned rats. We also compared the time spent exploring a conspecific during a given encounter between sham-control and STN-lesioned rats, using a two-sample Wilcoxon-Mann-Whitney test. For social and objects discrimination tests, we compared the exploration time of two stimuli during the second encounter using Wilcoxon-paired mean test, within each group. We also compared the time spent exploring a conspecific during the first encounter between sham-control and STN-lesioned rats using two-sample Wilcoxon-Mann-Whitney tests and between optogenetic groups using Kruskal-Wallis tests followed by Dunn multiple comparisons for post hoc analysis. Similarly, for the cage-mate recognition test, we compared the time spent exploring the cage-mate versus the unfamiliar rat, within group, using Wilcoxon-paired mean test.

Regarding the extracellular recordings of STN neurons activity, due to interindividual variability, we first normalized the data on the 20s of baseline acquisition for each cell. Data were further analyzed using Friedman tests, followed by Dunn multiple comparisons, as the variation compared to baseline activity (activity at t1 minus baseline activity, divided by baseline activity).

## Results

Representative intact and lesioned STN, as well as viral expression by EYFP-tracing fluorescence and optic fiber implantation, are illustrated in **Figure 1**. One rat was excluded from STN-lesioned group due to unsatisfying lesion (partially outside the STN). In the optogenetic groups, three rats were excluded for absence of fluorescence in either one or both STN, and three others for optic fiber misplacement.

**Figure 1:**
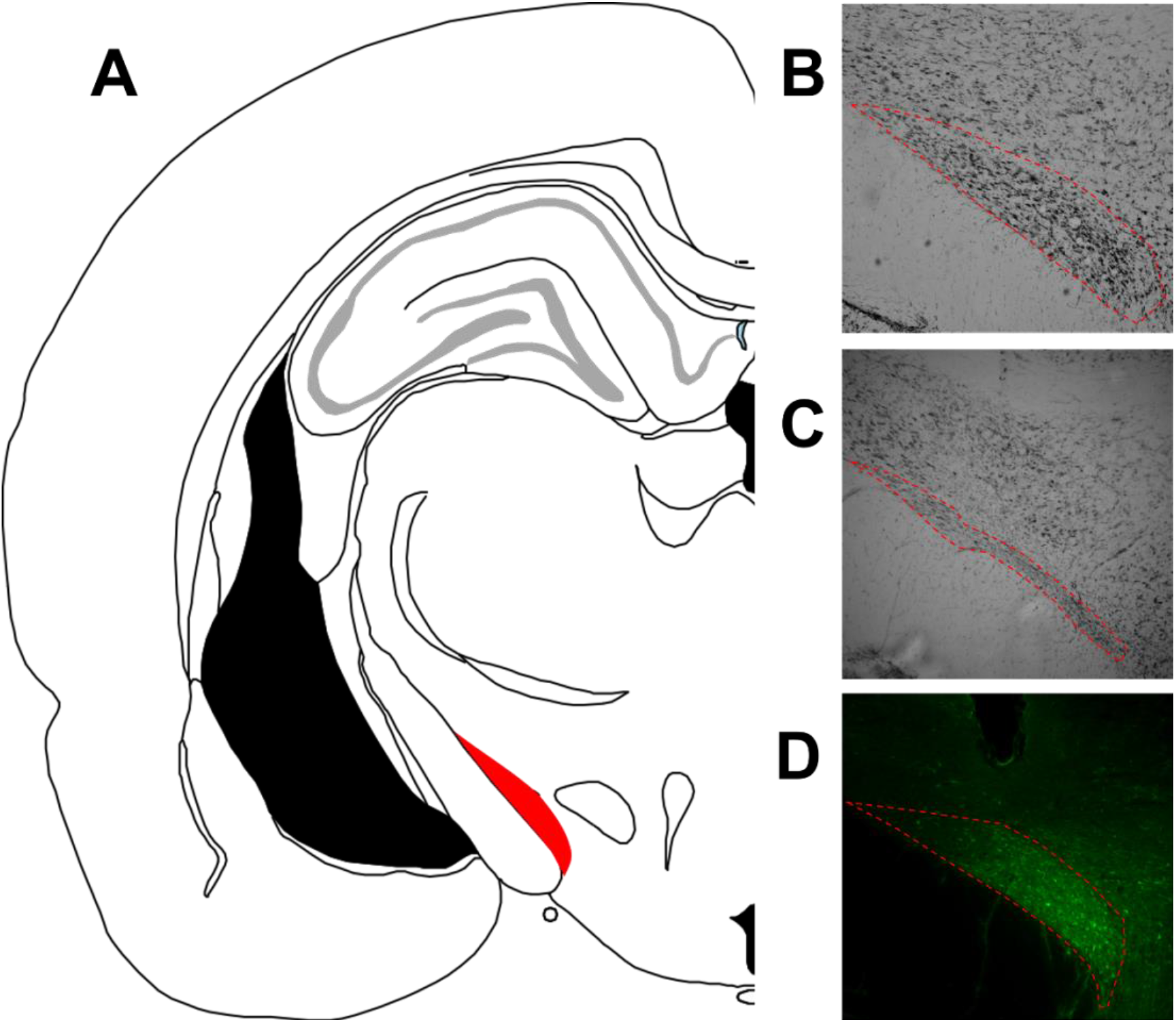
Frontal sections of the subthalamic nucleus. A: Schematic coronal section of the rat brain at the level of the STN (left STN in red), from Paxinos & Watson. Right panels: Representative frontal sections of the STN stained with thionine, with the STN delineated by the red lines in a (B) sham operated rat and (C) STN-lesioned rat. The lesions were characterized by neuronal loss and gliosis. Representative native fluorescence emitted by the transfected STN neurons and implanted optic fiber trace (D).

### STN lesions blunt social recognition in the Habituation-Dishabituation test

To examine a possible involvement of the STN in social recognition, we first employed a social habituation-dishabituation paradigm (**Figure 2**). Friedman test highlighted a significant effect of the encounters on the duration of exploration in sham-control rats (n=6) (χ^2^(4)=19.87, p=0.0005). During the habituation phase, post-doc analysis showed a continuous decrease in the amount of time spent at exploring the stimulus animal (encounter 1 vs 2: p=0.4014, vs 3: p=0.0076, vs 4: p=0.0002), suggesting that sham-control rats recognized this peer, while familiarity with the stimulus animal increases. The time spent to explore the novel conspecific during the dishabituation phase was significantly higher than during the fourth encounter of habituation (encounter 4 vs 5: p=0.0247), but not when compared to the first one (encounter 1 vs 5: p=0.8050). These results confirm that the decrease in exploratory behavior during the habituation phase results of prior encounter with the same conspecific and introduction of novelty restores exploration. In contrast, in the STN-lesioned animals (n=8), Friedman test showed no modulation in the amount of time spent at exploring the stimulus rat across encounters (χ^2^(4)=3.8, p=0.4337). Confirming this deleterious effect of STN lesions on peer recognition, during the fourth encounter of habituation, sham-control rats spent significantly less time exploring the ‘now-familiar’ conspecific than STN-lesioned rats (U=2, n=14, p=0.0027). On the other hand, sham-control and STN-lesioned animals showed the same amount of time exploring a new stranger conspecific during the first (U=19, p=0.5728) and five (U=20, p=0.6620) encounter, suggesting that STN-lesion does not affect social investigation of a stranger peer, according to Reymann et al. (2013).

**Figure 2:**
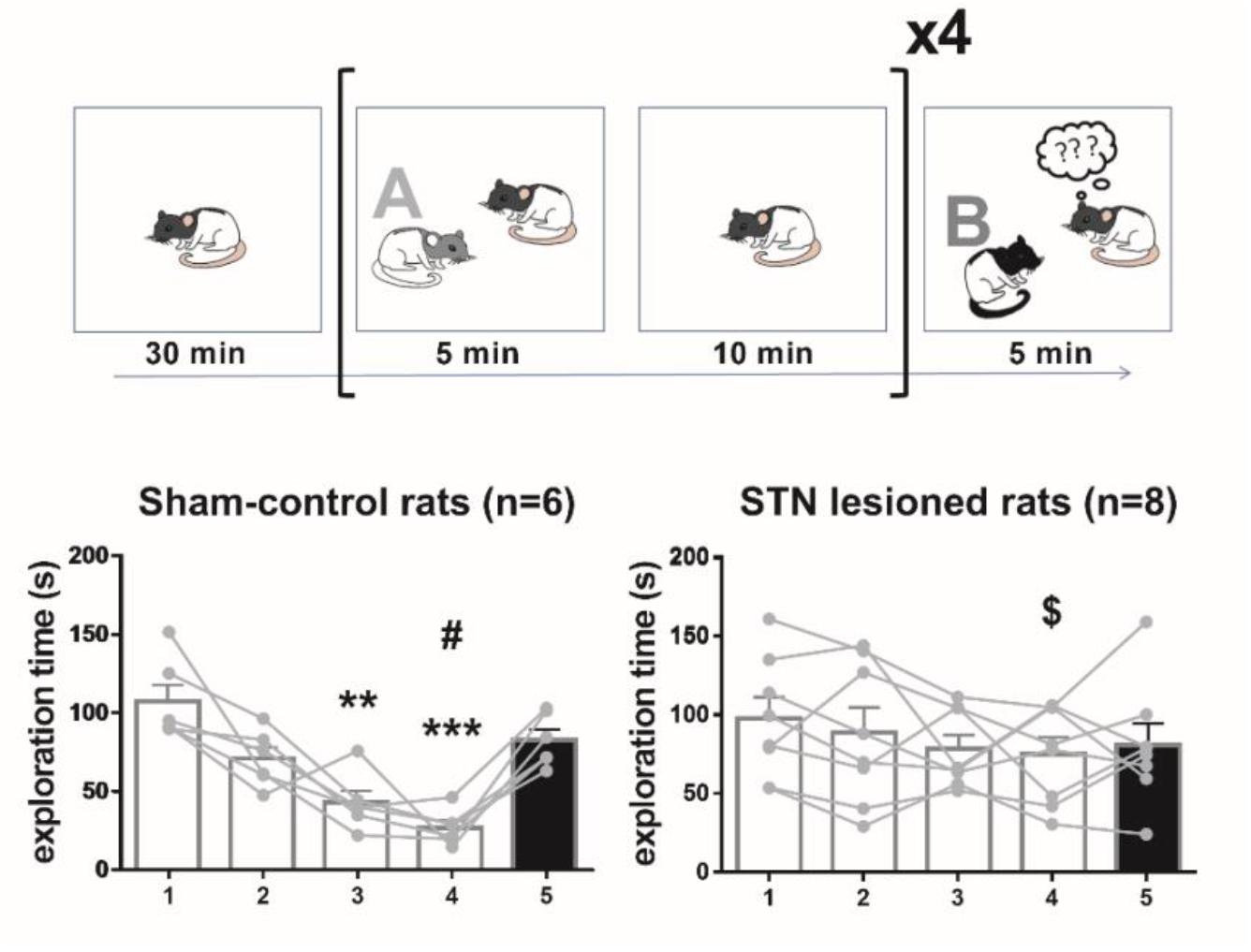
STN lesions affect social recognition in the Habituation-Dishabituation test. Upper part: Schematic representation of the procedure. After a 30-min habituation to the arena (Left), the subject rat was subjected the habituation phase (in the black brackets), consisting in 4 repetitions of the following sequence: a 5-min encounter to the stimulus animal A (in grey and white) and a 10-min interval (subject alone). Then, the subject was exposed for 5-min to a novel unfamiliar rat (B, in black), for the dishabituation phase (right). Lower part: Mean time (±SEM) spent by the sham-control (n=6, left) and STN-lesioned (n=8, right) rats exploring the stimulus animals (A white bars; B black bars) during each 5-min encounter. *\*\*: p<0*.*01 and ***: p<0*.*001 compared with encounter 1. #: p<0*.*05, compared with encounter 5. $: p<0*.*05, compared with sham-control rats*. The grey circles illustrate the individual data.

### STN neuromodulation affects social memory, but not immediate social discrimination

To verify and better characterize the involvement of STN on peer recognition, sham-control and STN-lesioned rats were subjected to two social discrimination tests, during which the 2^nd^ encounter took place immediately after the 1^st^ one (**Figure 3**), or after a 30-min intertrial interval during the two encounters (**Figure 4**). Independently of the intertrial delay, sham-control and STN-lesioned rats spent the same amount of time exploring the social stimulus during the 1^st^ encounter (sham control vs STN-lesioned rats without delay: U=19, p=0.5728 and with 30-min delay: U=20, p=0.6620). These results confirm that STN lesions do not affect social investigation. As illustrated in **Figure 3A**, when the 2^nd^ encounter immediately followed the 1^st^ encounter, both sham-control (W=21, p=0.0313) and STN-lesioned (W=36, p=0.0078) animals showed a significant preferential exploration for the novel rat over the familiar one, suggesting that STN lesions do not affect immediate peer discrimination, nor social novelty preference. However, following the 30-min intertrial interval (**Figure 4A**), while sham-control rats still showed a preferential exploration for the novel over the familiar animal (W=21, p=0.0313), the STN-lesioned rats spent the same time exploring the two social stimuli (W=0, p>0.9999), confirming that STN lesion impaired social recognition memory.

**Figure 3:**
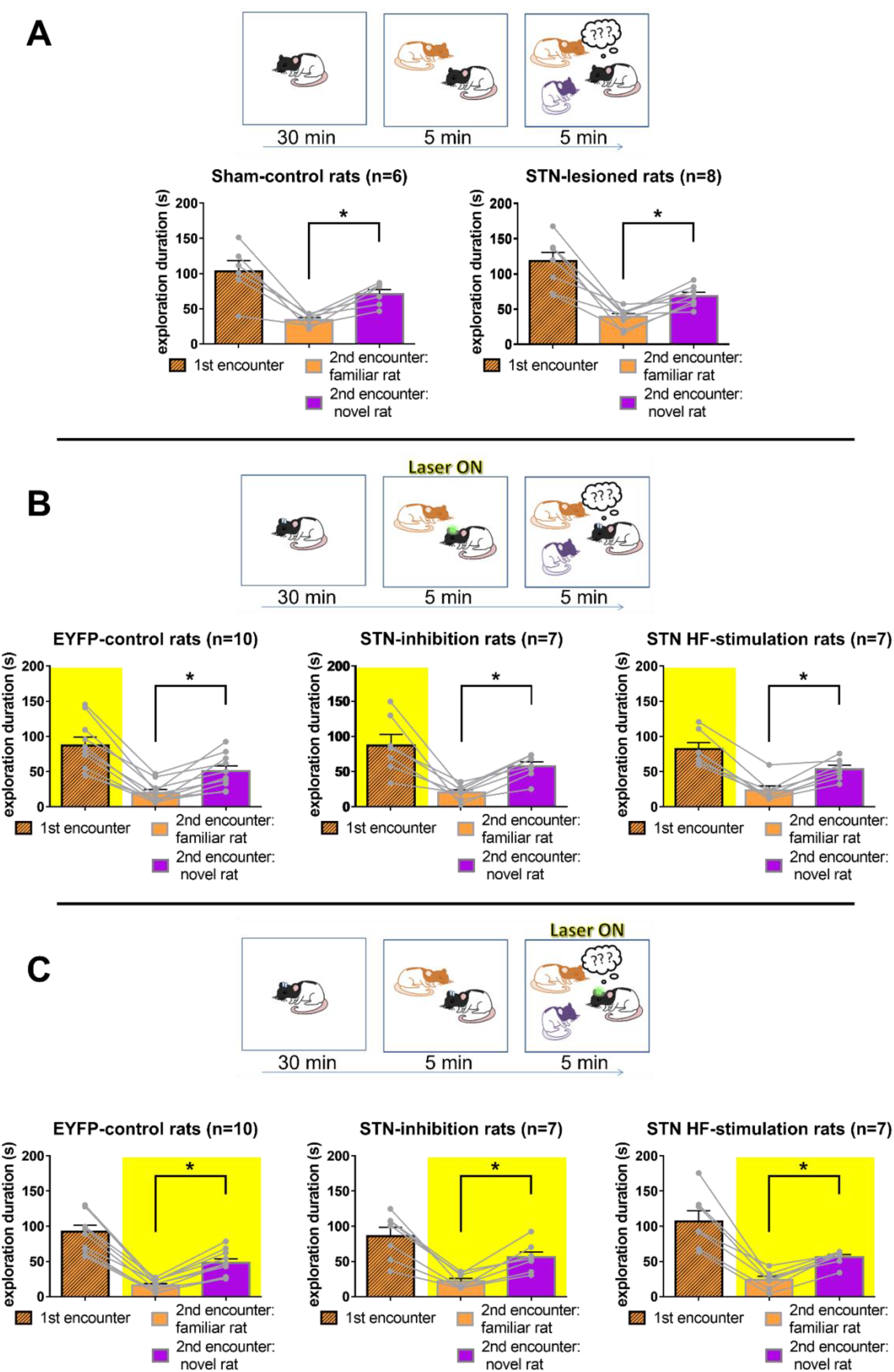
Neither STN lesions nor its optogenetic modulation affect immediate social discrimination. A: Effects of the STN lesions. Upper part: schematic representation of the procedure. After a 30-min habituation to the arena, the subject rat was exposed to a 5-min encounter with a stranger stimulus animal (1, in orange). At the end of the 5min, a new stranger stimulus rat (2, in violet) was introduced in the arena, for an extra 5-min encounter. Lower part: The graphs represent the mean exploration time (±SEM) spent by the Sham-control (n=6, left) and STN-lesioned (n=8, right). Dashed orange: exploration of stimulus rat 1 during the first encounter, orange: exploration of the stimulus rat 1 during the 2^nt^ encounter; Violet: exploration of stimulus rat 2. B and C Upper part: The rats with STN optogenetic manipulation have been subjected twice to this test, in a counter-balanced order: once, with the laser ON during the first, but not the second encounter (B), and once with the laser OFF during the first encounter and ON during the second one (C). B and C lower part: The graphs represent the mean exploration time (±SEM) spent by the subject rat of the EYFP-control (n=10, left), the STN inhibition (n=7, center) or the STN HF-stimulation (n=7, right) groups (B and C) exploring the stimulus animals 1 during the first encounter (dashed orange), the same stimulus animal 1 (orange) or the novel one (2, violet) during the 2^nd^ encounter. *\*:p<0*.*05 compared with the exploration duration of the novel stimulus rat*. The yellow area represents the period of activity of the laser. The grey circles illustrate the individual data.

**Figure 4:**
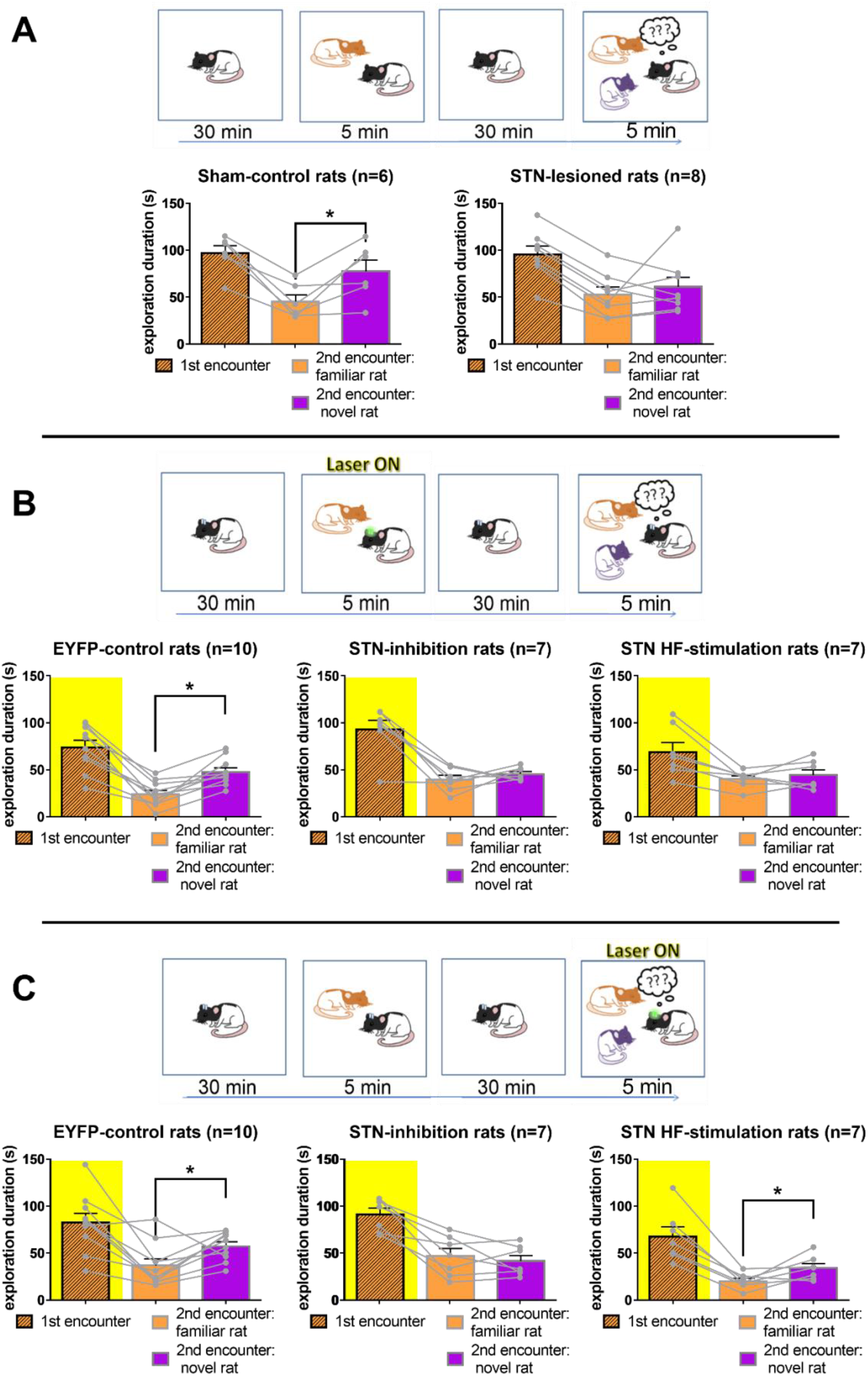
Both STN lesions and its optogenetic modulation alter social discrimination memory. Upper part of A, B, C: Schematic representation of the procedure. After a 30-min habituation to the arena, the subject rat was subjected to a 5-min encounter with an unfamiliar stimulus animal (1, in orange), constituting the encoding phase. After a 30-min intertrial interval, the subject animal was re-exposed for a 5-min recall encounter to the same stimulus animal 1, which became familiar to the subject, simultaneously with a novel unfamiliar stimulus animal (2, in violet). A relates to the procedure testing STN lesions. B and C: The rats subjected to STN optogenetic manipulations performed twice the test, in a counter-balanced order: once, with the laser ON during the 1^st^, but not the 2^nd^, encounter (B), and once with the laser OFF during the 1^st^ encounter and ON during the 2^nd^ one (C). Lower part of A, B, C: The graphs represent the mean exploration time (±SEM) spent by the Sham-control (n=6, left) and STN-lesioned (n=8, right) (A), or the EYFP-control (n=10, left), STN inhibition (n=7, center), STN HF-stimulation (n=7, right) rats 1 (B and C) exploring the stimulus animals 1 during the 1^st^ encounter (dashed orange), the same stimulus animal 1 (orange) or the novel stimulus rat 2 (violet) during the 2^nd^ encounter. *\*:p<0*.*05 compared with the exploration duration of the novel stimulus rat*. The yellow area represents the period of activity of the laser. The grey circles illustrate the individual data.

To explore a possible effect of STN manipulation on a subprocess of social discrimination memory, 10 EYFP-control 9 STN optogenetic inhibition, 7 STN optogenetic HF-stimulation rats were subjected to the same paradigms.

To first assess a possible effect of STN modulation on encoding of social information, optogenetic rats were subjected to the laser activation during the 1^st^ encounter. EYFP-control, STN inhibition and STN HF-frequency rats spent approximatively the same time exploring the stimulus animal during the 1^st^ encounter (without delay: χ^2^=0.2015, p=0.904, and with 30-min delay: χ^2^=3.783, p=0.1509), suggesting that STN optogenetic modulation does not alter social investigation. When the 2^nd^ encounter immediately followed the 1^st^ encounter, the 3 groups spent significantly more time exploring the novel over the familiar rat (EYFP-control: W=55, p=0.0020, STN inhibition: W=28, p=0.0156 and HF-stimulation rats: W=28, p=0.0156) (**Figure 3B**), showing that the photo-modulation of STN activity during the encoding phase does not alter peers discrimination, nor social novelty preference. However, following the 30-min delay, while EYFP-control rats still showed a preferential investigation of the novel over the familiar rat (W=55, p=0.0020), STN inhibition (W=7, p=0.5625) and HF-stimulation (W=5, p=0.6875) rats spent the same amount of time exploring both social stimuli (**Figure 4B**). Thus, both STN optogenetic inhibition and HF-stimulation, when performed during the encoding of social information, alter social discrimination memory.

To assess a possible effect of STN photo-modulation on social information recall, the same optogenetic rats were subjected to the laser activation during 2^nd^ encounter. The three groups spent approximatively the same time exploring the stimulus animal during the 1^st^ encounter (without delay: χ^2^=1.621, p=0.4447, and with 30-min delay: χ^2^=3.054, p=0.2172). When the 2^nd^ encounter immediately followed the 1^st^ encounter, the 3 groups spent significantly more time exploring the novel over the familiar stimulus rat (EYFP-control: W=55, p=0.0020, STN inhibition: W=28, p=0.0156 and STN high frequency stimulation rats: W=28, p=0.0156) (**figure 3C**), confirming that STN photo-modulation during social information recall does not alter immediate peer recognition, nor social novelty preference. However, following the 30-min intertrial delay (**Figure 4C**), while EYFP-control rats still showed a preferential investigation of the novel vs the familiar rat (W=39, p=0.0488), STN inhibition rats spent approximatively the same amount of time exploring both social stimuli (W=4, p=0.8125). Therefore, social memory discrimination is altered by STN optogenetic inhibition during the recall phase of social information. Interestingly, STN HF-stimulation rats also showed a preferential exploration of the novel over the familiar stimulus (W=24, p=0.0469), revealing that STN optogenetic HF-stimulation during the recall phase does not alter social discrimination memory.

Altogether, these results suggest that STN lesions affect social memory recognition. And while STN optogenetic inhibition alters social discrimination memory when performed during both encoding or recall, its optogenetic HF-stimulation suppresses social discrimination memory only when performed during the encoding, but not recall, phase.

### STN neuromodulation does not affect object recognition memory

These previous results might rely on the involvement of the STN in general memory processes. To further explore this possibility, all rats were subjected to an object discrimination memory paradigm, as illustrated **in Figure 5**. In this paradigm, both sham-control (W=21, p=0.0313) and STN-lesioned rats (W=32, p=0.0234) showed a preferential exploration of the novel over the familiar object during the 2^nd^ exposure (**Figure 5A**). When the laser was ON during the 1^st^ exposure, both EYFP-control (W=41, p=0.0371) and STN inhibition rats (W=24, p=0.0469) spent more time exploring the novel vs the familiar object during the 2^nd^ exposure. (**Figure 5B**) Results were less obvious for the STN HF-stimulation group (W=18, p=0.1563). It is noteworthy that only one STN HF-stimulation showed a strong preference for the familiar over the novel object (red line in the figure). If this rat’s scores were removed from the sample, the time spent exploring the novel object was significantly higher over the familiar for the rest of the group (W=21, p=0.0313). Nevertheless, we report here the result from the full sample as there is no reason to believe that this outlier rat results should be removed from analysis. Similarly, when the laser activation took place during the 2^nd^ exposure, both EYFP-control (W=55, p=0.0020) and STN HF-stimulation group (W=24, p=0.0469) showed a preferential exploration for the novel object over the familiar one (**Figure 5C**). For the STN inhibition rats, results were less obvious (W=16, p=0.2188). Once again, this result was highly impacted by the preference of only one rat for the familiar over the novel object (red line in the figure). If we removed this rat from the sample, STN inhibition rats’ preference for the novel over the familiar object appeared to be significant (W=21, p=0.0313).

**Figure 5:**
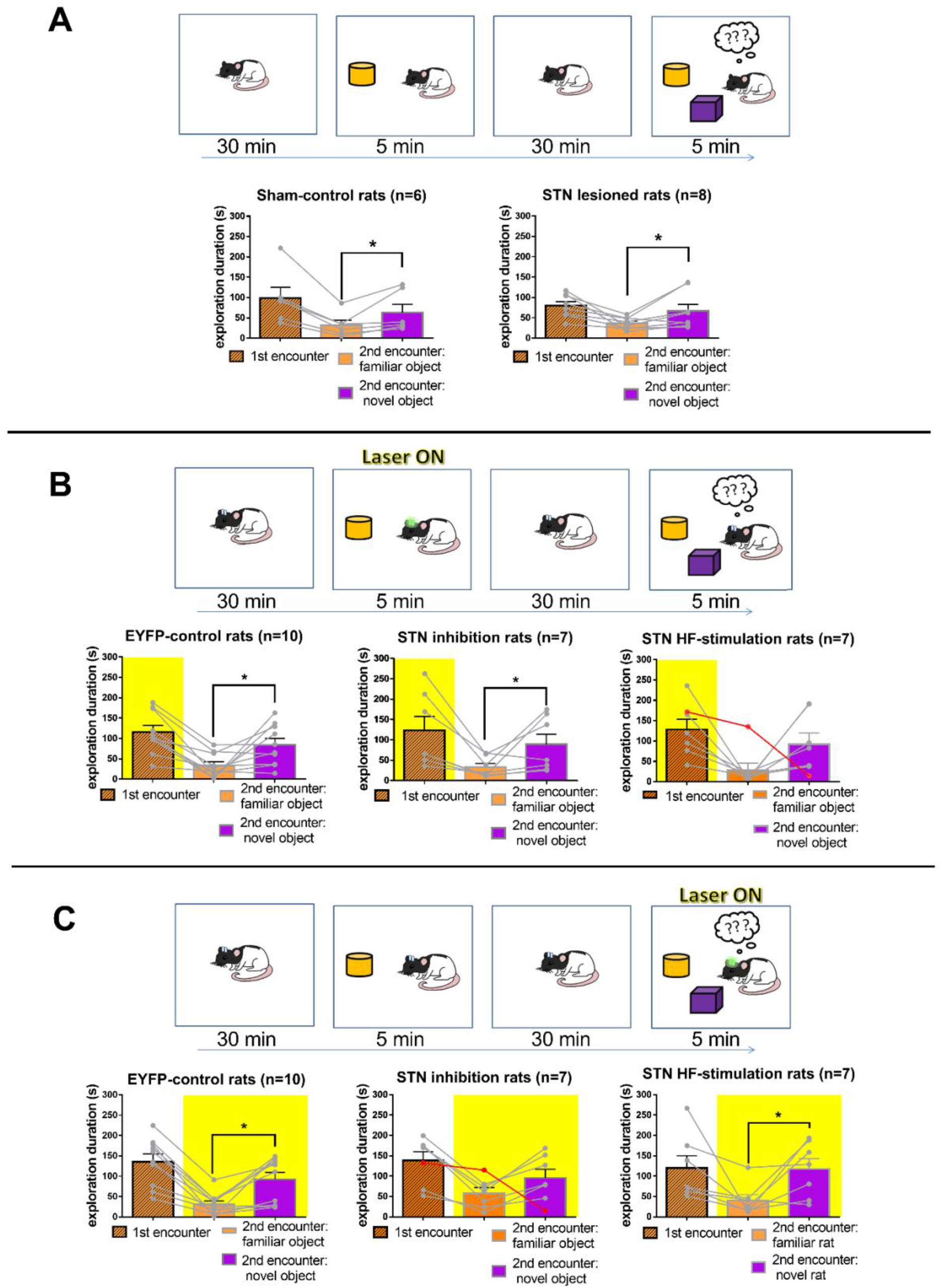
Neither STN lesions nor its photo-modulation seem to alter objects discrimination memory. Upper part A, B and C: Schematic representation of the procedure. After a 30-min habituation to the arena, the subject rat was subjected to a 5-min exposure with non-familiar stimulus object (1, in orange), constituting the encoding phase. After a 30-min intertrial interval, the subject rat was re-exposed for 5-min to the same stimulus object 1, which became familiar to the subject, simultaneously with a novel unfamiliar stimulus object (2, in violet). Panel A relates to the experiment testing STN lesions. B and C: The rats with STN optogenetic manipulations were subjected twice to this test, in a counter-balanced order: once, with the laser ON during the 1^st^, but not the 2^nd^ exposure (B), and once with the laser OFF during the 1^st^ encounter and ON during the 2^nd^ one (C). Lower part A, B, C: The graphs represent the mean exploration time (±SEM) spent by the Sham-control (n=6, left) and STN-lesioned (n=8, right) (A), or the EYFP-control (n=10, left), the STN inhibition (n=7, center) and the STN HF-stimulation (n=7, right) rats (B and C) exploring the stimulus objects during the 1^st^ exposure (dashed orange), the same object (orange) and the novel one (violet) during the 2^nd^ exposure. *\*:p<0*.*05 compared with the exploration duration of the novel object*. The yellow area represents the period of activity of the laser. The grey circles illustrate the individual data. The red line and dots correspond to the results of the outlier animals.

Altogether, these results highlighted the specific involvement of the STN neuromodulation in peer recognition memory. We showed that STN is not involved in general memory, nor in novelty preference, in line with Pelloux et al., (2014).

### Optogenetic inhibition, but neither HF-stimulation, nor lesion of the STN impairs cage-mate discrimination memory

To explore the extent of the alteration of social memory induced by STN neuromodulations, sham-control, STN-lesioned and optogenetic rats were subjected to a cage-mate discrimination memory test, as illustrated in **Figure 6**. Both sham-control (W=21, p=0.0313) and STN-lesioned (W=28, p=0.0156) rats showed a significant preferential exploration of the stranger rat over their cage-mate (**Figure 6A**). Thus, the STN lesions did not affect the cage-mate discrimination memory. Similarly, both EYFP-control (W=53, p=0.0039) and STN HF-stimulation (W=24, p=0.0469) groups spent significantly more time exploring the stranger rat than their cage-mate (**Figure 6B**). However, STN-inhibition rats spent approximately the same amount of time exploring the two stimulus animals (W=2, p=0.9375). Thus, STN optogenetic inhibition, but neither its HF-stimulation nor its lesions, impaired the cage-mate discrimination memory.

**Figure 6:**
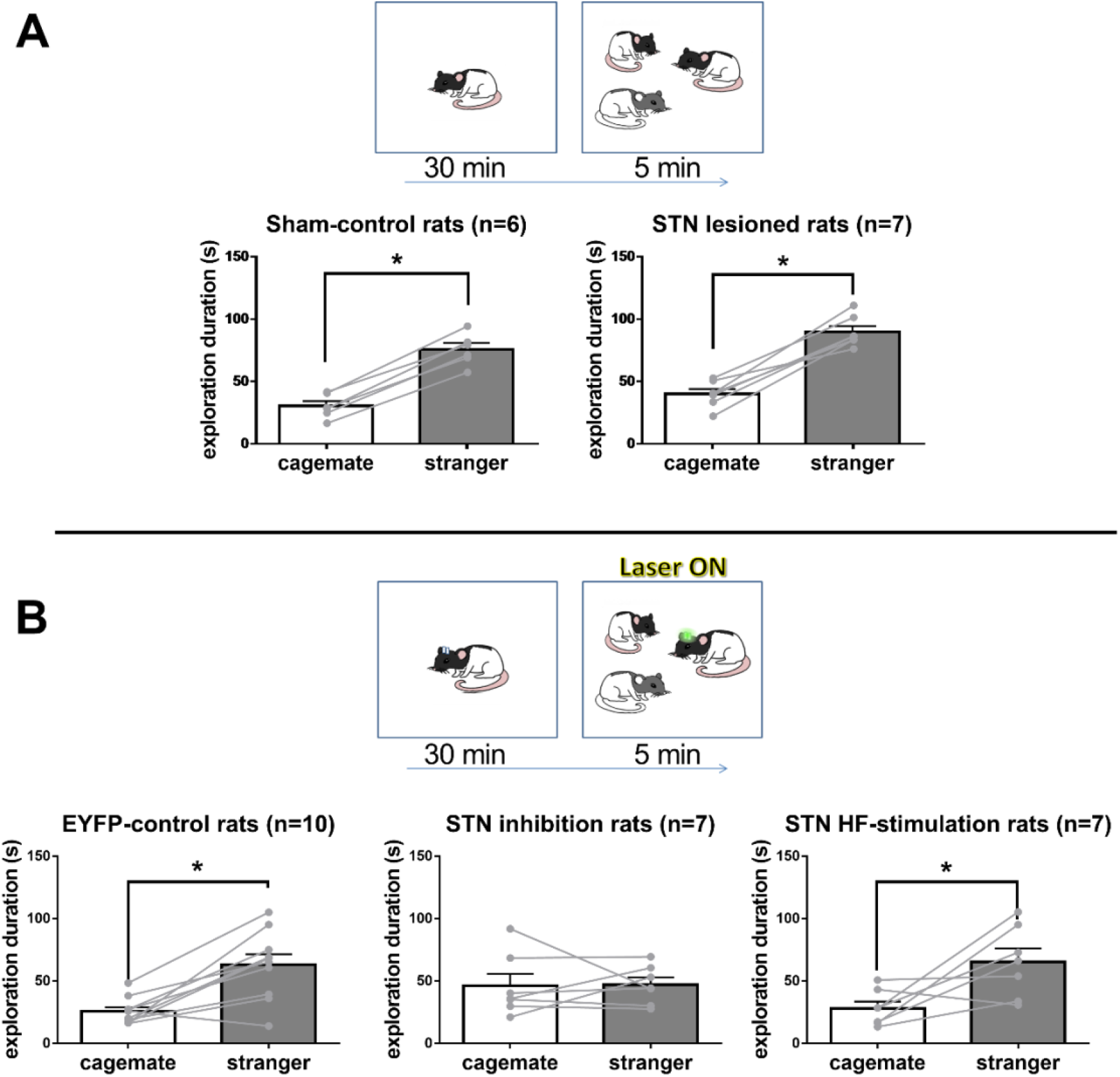
STN optogenetic inhibition, but neither its HF-stimulation nor its lesions, alters rats’ ability to recognize their cage-mate. Upper part A and B: Schematic representation of the procedure. After a 30-min habituation to the arena, the subject rat was subjected to a 5-min encounter with both its cage-mate and a stranger stimulus rat. A relates to the experiment testing STN lesions, B refers to the experiment testing optogenetic manipulations. Lower part: The graphs represent the mean exploration time (±SEM) spent by the Sham-control (n=6, left) and STN-lesioned (n=7, right) (A), or the EYFP-control (n=10, left), the STN inhibition (n=7, center) or the STN HF-stimulation (n=7, right) rats (B) exploring the cage-mate (white bars) or the stranger stimulus rat (grey bars). *\*:p<0*.*05 compared with the exploration duration of the stranger stimulus rat*. The grey circles illustrate the individual data.

### *In vivo* electrophysiological assessment of STN optogenetic inhibition and HF-stimulation

To assess the efficacy of our optogenetic manipulations on STN neuronal activity, we performed in vivo recordings of STN activity in anesthetized animals naïve to the behavioral experiment (**Figure 7**). Rats’ STN were injected with virus expressing the opsin ARCHT3.0 (n=3, STN inhibition) or hChR2(E123T/T159C) (n=7, STN HF-stimulation). Animals were anesthetized before being subjected to STN photo-modulation. The activity of STN cells was recorded during 20s of baseline activity (laser OFF), 180s of laser bins (i.e., 9 consecutive alternations of light stimulation at 5mW, interleaved with 5s periods of obscurity for STN inhibition or continuous 130Hz light stimulation for STN HF-stimulation) and 20s post laser stimulation (laser OFF). To estimate the population response, the neuronal firing of STN neurons was normalized to the 20s baseline prior STN photo-modulation.

**Figure 7:**
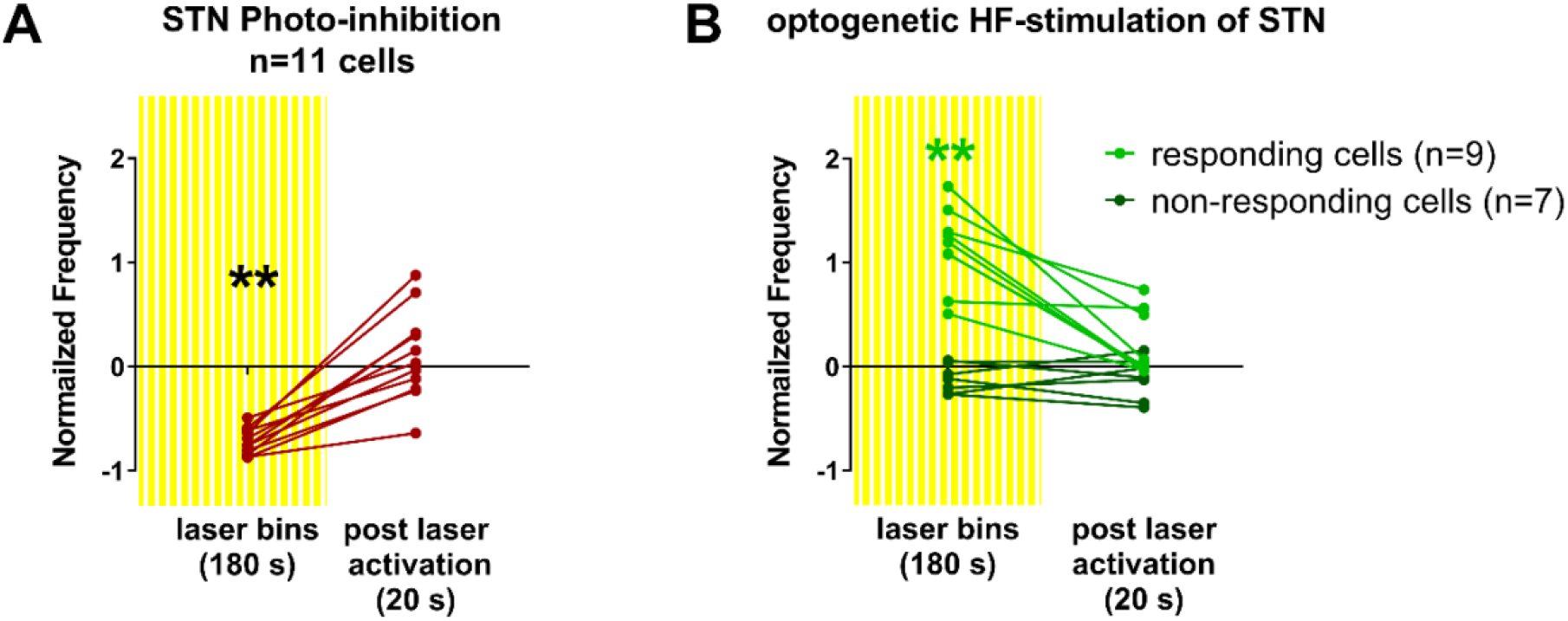
*In vivo* electrophysiological assessment of STN optogenetic inhibition and HF-stimulation. Frequency of firing rate in STN neurons, normalized on the 20s of baseline activity, then subjected to 180s of laser bins activation (dashed yellow), consisting in alternations of 15s of lights pulse interleaved by 5s period of obscurity for STN inhibition (A, n=11 cells) and continuous 130Hz light pulse for STN HF-stimulation (B, n=9 non responding cells in light green and n=7 non-responding cells in dark green), and the followed 20s post laser activation. *\*\*:p<0*.*0001, compared with baseline (only in responding cells in B)*.

Friedman test revealed a significant effect of these 3 conditions on the neuronal activity (χ^2^(2)=16.55, p<0.0001) for the 11 cells recorded in animals subjected to STN photo-inhibition (**Figure 7A**). Compared with the baseline activity, Dunn’s post-hoc comparisons confirmed that neuronal activity was significantly reduced (p=0.0013) by the laser bins activation compared with the baseline activity. Post laser activation, cells returned to a baseline level of activity (p>0.9999).

Interestingly, only 9/16 of the cells recorded in STN HF-stimulation rats responded to the laser activation (**Figure 7B**; ‘responding cells’). In these cells, Friedman test showed a significant effect of the conditions of laser activation (χ^2^(2)=13.56, p=0.0003). Dunn’s post-hoc comparisons revealed that neuronal activity increased during the 180s of laser bins activation (p=0.0019) before returning to a baseline level of activity (p>0.9999). In contrast, Friedman test show no effect of the conditions of laser activation in the 7 non-responding cells (χ^2^(2)=2, p=0.4861).

## Discussion

Here, we showed that STN-lesioned rats failed to recognize an previously encountered peer in the habituation-dishabituation paradigm and to discriminate between a familiar and a novel conspecific in the social discrimination test with 30-min intertrial between the two encounters. These results cannot be explained by a general memory impairment, since STN-lesioned rats showed the same ability to discriminate between a familiar and a novel unfamiliar object than sham-control rats. Furthermore, the fact that STN-lesioned and sham-control rats spent similar amount of time exploring a social stimulus during the 1st encounter in the habituation-dishabituation and social discrimination tests reveal that the STN lesion does not alter social exploration, in line with Reymann et al., (2013). Moreover, in the social discrimination test without delay between two encounters, STN-lesioned rats are as able as sham-control rats to discriminate between the novel and the previously encountered peer, indicating that the STN does not influence social novelty preference, nor immediate peer recognition. Finally, STN-lesioned rats showed the same ability than sham-control ones to recognize their cage-mate. Altogether, our results revealed that STN lesion specifically blunts rat social recognition memory for a recently encountered peer.

This study is the first, to our knowledge, illustrating the STN as a key component of social recognition memory. It has been shown that the presence of a peer and the playback of positive USV are both rewarding, and their familiarity further modulate their rewarding properties (Giorla et al., 2022; Vielle et al., 2021). Indeed, these social stimuli appeared to be more rewarding if they are stranger (i.e., presence of an unfamiliar rat or USV emitted by an unfamiliar rat) than familiar with the subject (i.e., presence of the cage-mate or USV emitted by the cage-mate). Interestingly, in these studies, STN lesions abolished the familiarity-related modulation of the social stimuli rewarding value. Considering that the STN is involved in social recognition memory, it would be possible that its lesions altered rats’ ability to recognize their cage-mate, resulting in an undifferentiated value of stranger *vs*. familiar social stimuli. Although in our test, STN-lesioned rats were able to recognize their cage-mate, it does not mean that it was also the case in these previous studies. Indeed, in Vielle et al. (2021), rats only had access to the USV emitted by their peers, which could be insufficient to allow STN-lesioned rats to evaluate the familiarity of the USV emitter. Similarly, in Giorla et al. (2022), rats were separated from social stimuli by a grid during cocaine self-administration and social stimuli were encaged during the social preference test of a stranger or familiar rat over an object. Making measurements easier, this practice is widely used in social cognition and preference literature (see for instance Jacobs & Tsien, 2017; Lukas et al., 2011; Tan et al., 2019). However, separation of the compartments containing social stimuli prevents the rats from having access to the nonvolatile fraction of the individual’s olfactory signature (via the body surface and notably the anogenital region) (Engelmann et al., 2011). With a blunted social recognition memory and deprived from these sensorial indices, STN-lesioned rats would be unable to recognize their cage-mates. On the contrary, in our test, rats could freely explore the social stimuli and then use all sensorial indices to discriminate their cage-mates from unfamiliar rats, allowing a proper recognition performance of the STN-lesioned rats.

Overall, the implication of the STN in social recognition memory may have critical consequences for the follow-up of patients treated with STN-DBS for PD or OCD. If social behavioral maladjustments are frequently reported in PD patients following STN-DBS (Houeto et al., 2002; Perozzo et al., 2001), as well as deficits in facial and vocal emotions reading (Biseul et al., 2005; Brück et al., 2011; Kalampokini et al., 2020; Péron, Grandjean, et al., 2010), faces recognition seems to be preserved (Biseul et al., 2005; Drapier et al., 2008; Péron, Biseul, et al., 2010). Whether it is a discrepancy with rat data, or the lack of deficits reported in PD patients is due to differences in experimental paradigms remains to be determined. The sensorial modality tested could play a role in the possible deficits observed. Indeed, faces recognition only recruits visual modality, contrary to rodent recognition paradigms presented here. Does it mean that human social brain only processes visual information? Contradicting this assertion, recent literature has revealed the essential, while unconscious, contribution of the olfactory system in human cognition (Endevelt-Shapira et al., 2018; Frumin et al., 2015; McGann, 2017). Alternatively, these discrepancies between results obtained from humans and rats could rely on the type of social recognition memory which is tested. Tranel & Damasio (Tranel & Damasio, 1987, 1988, 1990, 1993) describe the overt (i.e., self-reported, conscious) *vs*. covert (i.e., implicit, unconscious) forms of social memory, mediated by two different neural networks. Their work revealed that the covert form of learning is driven by affective salience. Rats’ performance in social recognition tests being driven by affective salience (i.e., novelty-reinforcing) (Engelmann et al., 1995; Thor & Holloway, 1982), they reflect covert learning. In contrast, clinical studies generally explore the overt form of social memory (i.e., declarative), directly questioning the patient’s ability to recognize an individual. Considering the role of the STN in incentive salience and in emotion recognition (Pelloux et al., 2014; Péron et al., 2013; Serranová et al., 2011), the STN appears as a serious candidate for such covert affective-driven social memory processes. Overall, further studies are needed to fully understand the role of the STN in social recognition memory in human and the relative consequences of STN DBS.

To better characterize the involvement of the STN on social recognition memory, we used an optogenetic approach, allowing us to stimulate at 130 Hz or to inhibit STN neuronal activity specifically during the encoding or recall phase of social memory. Like its lesions, STN optogenetic modulations did not to affect social novelty preference, immediate peer recognition, nor general social exploration. Furthermore, STN optogenetic modulations did not seem to alter object memory. Yet, our results from the peer discrimination memory (i.e., with 30-min delay between the two encounters) show that STN optogenetic inhibition alters social recognition, when it is performed during both encoding or recall phase. In contrast, STN HF-stimulation alters rats’ ability to recognize a peer, only when applied during the 1^st^, but not the 2^nd^ encounter. Furthermore, results from the cage-mate recognition showed that STN HF-stimulation group discriminated their cage-mate from a stranger peer after 30-min of separation, confirming that STN HF-stimulation does not alter social recognition memory, when performed during the recall phase.

The fact that STN short-term (optogenetic), but not long term (lesions), inhibition blunts rats’ ability to recognize their cage-mate suggest a compensatory mechanism. For instance, STN modulation could affect social olfaction recognition, a well-known highly salient sensory modality for rodents. In this hypothetical scenario, STN-lesioned rats used to interact with their cage-mates with this olfactory deficit for weeks since STN lesioning, which is not the case for rats subjected to STN optogenetic inhibition. STN-lesioned probably rely on others sensory cues, as a compensatory mechanism, to recognize their cage-mate. In contrast, for optogenetic rats, the sudden alteration of their olfactory abilities induced by the laser activation during the test would be disabling to recognize their cage-mate. Further investigations are necessary to confirm and better characterize this compensatory mechanism of the STN lesions.

When performed during the 1^st^ encounter of the social discrimination memory test, both optogenetic manipulations blunted social recognition. This similar behavioral effect of two opposite STN modulations has already been observed in previous study (Tiran-Cappello et al., 2018). It could rely on this assertion from Parent and Hazrati (1995): ‘the subthalamic nucleus seems to process in parallel cortical information of different types and convey them to the basal ganglia along separate channels’. As such, if an output structure needs to receive information from both the STN and the cortex, in parallel, then any modulation of the STN activity (i.e., stimulation and inhibition) could disrupt this circuitry and lead to a misinformation of this output structure. Then the STN could be represented as an integration relay necessary to process cortical information. Interestingly, Péron et al. (2013) also proposed a model stating that the STN produces a neural co-activation, temporally organized, in both cortical and subcortical levels, which is necessary to generate emotions and related feelings. Alternatively, this similar behavioral result of opposite optogenetic manipulations of the STN would rely on a more complex internal organization of this structure than what is known today (see for instance Wallén-Mackenzie et al., 2020). Further studies are necessary to elucidate this anatomical–functional organization of the STN, and its complex roles in behavior.

When performed during the 1^st^ encounter of the social discrimination memory test, both optogenetic manipulations blunted social recognition. The fact that similar behavioral effects result from two opposite neuromodulations is surprising. While not all the cells responded to the laser activation, the population response confirmed the efficiency of both optogenetic manipulations to inhibit or stimulate neural activity. However, the similarity in behavioral effects induced by both manipulations might be due to their blockade of local oscillatory activity. From a functional point of view, cerebral oscillatory activity would temporally coordinate neuronal ensembles in local and distal brain areas, inducing synaptic plasticity, thereby allowing information processing (acquisition, consolidation and combination of learned information) (Buzsáki & Draguhn, 2004). The presentation of an unfamiliar rat associated with an increased in alpha/theta band power and phase synchronicity respectively within and between limbic territories (medial and anterior olfactory bulb, medial amygdala, lateral septum and piriform cortex) (Tendler & Wagner, 2015). These changes in alpha/theta activity were further gradually normalized across repeated exposures with the same individual, suggesting that such oscillatory activity is driven by social novelty and contribute to social recognition memory. Interestingly, increase in theta power within the STN was associated with the affective-salience attribution to non-social rewards in PD. It is thus possible that the attribution of affective salience required for covert social memory is achieved by the development of theta activity within the STN. Any blockade of theta activity in the STN, through its lesion, optogenetic inhibition or HF-stimulation would then blunt the acquisition of social memory. Once social information is correctly encoded in the STN, through theta activity, any STN re-activation would be sufficient to recall it. Therefore, the recall of social information would by blunt by its inhibition (lesions or optogenetic inhibition), but not by its HF-stimulation. Further studies are needed to test this hypothesis, like the recording of STN oscillations during social memory encoding and recall.

## Conclusions

In conclusion, we found that the STN is specifically involved in social recognition memory. Both its optogenetic inhibition and its excitotoxic lesions impaired social recognition memory. STN optogenetic HF-stimulation leads to an alteration of social memory, only when performed during encoding, but not recall, of social information. None of these manipulations seem to interfere with social investigation, objects recognition memory, nor social novelty preference. STN optogenetic inhibition, but neither its HF-stimulation, nor lesions, even leads to an alteration of the cage-mate recognition memory. This differential effect of the STN lesions and optogenetic inhibition on the cage-mate recognition would rely on a compensatory mechanism. Further studies are needed to better characterize the role of the STN in social cognition, its place in the social brain circuitry and the possible consequences for the follow-up of patients treated with STN DBS these processes.

## Acknowledgements

Thanks to Alix Tiran-Cappello for his help with optogenetics tool and analyses and to Lucie Vignal for rats’ illustrations.

